# DNA methylation differences at regulatory elements are associated with the cancer risk factor age in normal breast tissue

**DOI:** 10.1101/101287

**Authors:** Kevin C. Johnson, E. Andres Houseman, Jessica E. King, Brock C. Christensen

## Abstract

**Background:** The underlying biological mechanisms through which epidemiologically defined breast cancer risk factors contribute to disease risk remain poorly understood. Identification of the molecular changes associated with cancer risk factors in normal tissues may aid in determining the earliest events of carcinogenesis and informing cancer prevention strategies.

**Results:** Here we investigated the impact cancer risk factors have on the normal breast epigenome by analyzing DNA methylation genome-wide (Infinium 450K array) in cancer-free women from the Susan G. Komen Tissue Bank (n = 100). We tested the relation of established breast cancer risk factors: age, body mass index, parity, and family history of disease with DNA methylation adjusting for potential variation in cell-type proportions. We identified 787 CpG sites that demonstrated significant associations (Q-value < 0.01) with subject age. Notably, DNA methylation was not strongly associated with the other evaluated breast cancer risk factors. Age-related DNA methylation changes are primarily increases in methylation enriched at breast epithelial cell enhancer regions (P = 7.1E-20), and binding sites of chromatin remodelers (MYC and CTCF). We validated the age-related associations in two independent populations of normal breast tissue (n = 18) and normal-adjacent to tumor tissue (n = 97). The genomic regions classified as age-related were more likely to be regions altered in cancer in both pre-invasive (n = 40, P=3.0E-03) and invasive breast tumors (n = 731, P=1.1E-13).

**Conclusions:** DNA methylation changes with age occur at regulatory regions, and are further exacerbated in cancer suggesting that age influences breast cancer risk in part through its contribution to epigenetic dysregulation in normal breast tissue.

## Background

Breast cancer represents a major public health problem in the United States with 245,000 new cases and 40,000 deaths expected this year [1]. An effective way to decrease disease-related morbidity and mortality is to identify individuals who may be at increased risk of developing breast cancer and apply early intervention strategies. In addition to inherited gene mutations, there are several demographic factors that are associated with an increased risk of breast cancer including: increasing age, being overweight after menopause, alcohol intake, having never been pregnant (that is, nulliparous), earlier age at menarche, and a family history of breast cancer [2–4]. However, the underlying biologic mechanism(s) through which many of these epidemiologically defined breast cancer risk factors contribute to carcinogenesis remains unclear.

Biomarkers strongly associated with breast cancer risk factors provide an opportunity to understand cancer development. One such potential biomarker investigated for its role in the early detection of breast cancer is DNA methylation. DNA methylation is the covalent addition of a methyl group to cytosine, often in the context of a cytosine followed by a guanine in the ‘5 to 3’ direction (that is, a CpG), and is necessary for cell-type specific differentiation, including in the mammary gland [5–7]. DNA methylation is a stable, yet modifiable epigenetic modification and DNA methylation alterations are known to occur early in breast carcinogenesis [8, 9]. It has been hypothesized that disease risk factors may mediate their disease predisposing effects through perturbation of epigenomic control. Candidate gene studies in tumor adjacent normal breast tissues indicate that DNA methylation changes are related to age as well as other known breast cancer risk factors. For example, women without breast cancer, but at a high risk (Gail model score) were more likely to have aberrant methylation of the tumor suppressor genes *APC* as well as *RASSF1* compared with low risk women [10]. In another candidate gene study of normal breast tissue, the same group observed that *RASSF1* methylation was associated with breast cancer risk level, and that increasing parity was associated with decreased *APC* methylation [11]. More recently, a study identified cancer-related field defects in DNA methylation using normal breast tissues from disease-free subjects and tumor-adjacent normal breast tissues Preliminary analyses in tumor-adjacent normal breast tissue provide evidence that age-related DNA methylation changes are more likely to be altered in breast tumors than randomly selected regions [13]. However, the relation of breast cancer risk factors with DNA methylation changes in the normal breast remains to be investigated.

Here we extend the foundational work to tissues from disease-free women with detailed breast cancer risk factor data and apply more comprehensive epigenomic profiling methods. We test the relation of breast cancer risk factors such as age, body mass index (BMI), reproductive, and family history with DNA methylation patterns using an epigenome-wide association study (EWAS) approach. Importantly, we include adjustment for potential variation in cellular proportions across samples. Age is the strongest risk factor for breast cancer and we have shown that the patterns of age-related DNA methylation are dependent upon genomic context and that these age-related methylation patterns were consistent across independent populations of normal breast tissue. We found that these molecular alterations become further altered in pre-invasive as well invasive cancerous lesions. Together, the epigenetic changes we identified here provide insights into how breast cancer risk factors are related to disease development.

## Results

### Differential DNA methylation is associated with breast cancer risk factors in normal breast tissues

Patient demographics and characteristics are presented in **Table 1**. The study participants ranged in age from 18 to 82 with a median age of 37. A small proportion of participants were underweight (2%; body mass index (BMI) < 18), 40% in the normal BMI range (>=18 and < 25), 30% were overweight (>=25 and < 30), and 28% were obese (>30). Over half of subjects had at least one full-term birth (56%), and the remaining 44% were nulliparous. To test the hypothesis that DNA methylation differences in normal breast tissue are related with known breast cancer risk factors we used the approach outlined in **Additional File 1**. Differences in cellular composition across samples represents a potential confounder when testing associations between DNA methylation and quantitative traits in epigenome-wide association studies (EWAS) Cellular proportions for each sample can be estimated through cytometric methods or by applying cell mixture deconvolution algorithms to DNA methylation measurement [15], [16]. Cellular proportions can then be incorporated into a statistical model as covariates to adjust for potential cellular heterogeneity. In the absence of direct cell counts or tissue-specific reference DNA methylomes, statistical methods that account for cell proportion variability across tissue samples without a reference DNA methylome have been widely used [15], [17–20]. To this end, we applied a convex non-negative matrix factorization approach to estimate the proportion of putative cell types in each tissue sample [20]. This approach provides an estimate of cellular proportions across a range of putative cell types (K). We identified the optimal number of putative cell-types as K = 6 as this estimate minimized the deviance of the bootstraps (see Methods, **Additional File 2A**). To investigate whether the heterogeneity in cellular proportions across samples were associated with phenotypic variables (e.g., subject age) we applied a quasi-binomial model for each subject. To avoid dependence on the selection of K (putative cell types) we examined associations over a range of evaluated K using a permutation test (1000 permutations) for inference of each phenotypic variable. As shown in **Additional File 2B**, estimated cell mixture proportions were significantly associated with subject age (Permutation *P*-value = 2.0E-03), but not subject BMI or parity (**Additional File 2B**).

**Table 1.**
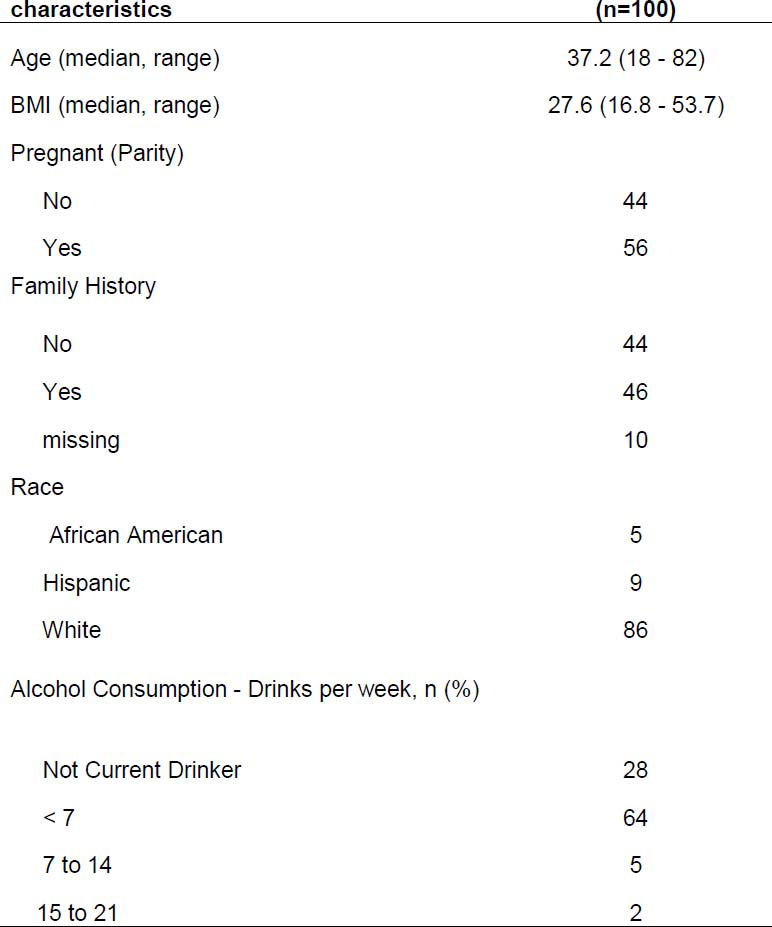
Subject demographics and characteristics (n=100)

To study the relation of DNA methylation with breast cancer risk factors we applied both unadjusted and cell-type adjusted linear models for microarray (*limma*) to examine the influence of subject age, BMI, and parity on the DNA methylome. Since the estimated cellular proportions for each sample sum to nearly one we include all but the estimated cell-type with the smallest proportion to avoid multi-collinearity in our models. In a multivariable limma model adjusted for differences in cellular mixtures 787 CpG sites were significantly associated with age, 0 CpG sites with BMI, and 0 CpG sites with parity after correcting for multiple hypothesis testing (*Q* < 0.01, **Figure 1A**). The full list of 787 CpG sites with genome annotation and statistical results is presented in **Additional File 3**. Notably, age-related DNA methylation alterations were predominantly hypermethylation events, that is, increased DNA methylation was associated with increased age (545 CpG sites, 69.3%). To assess the impact adjusting for cellular proportions had on the identification of significant associations and effect sizes we computed the difference between the coefficients (that is, a delta coefficient value) at each CpG for the models unadjusted and adjusted for cell-type. A large CpG-specific delta value provides evidence for associations between DNA methylation and risk factors that may be most confounded by differences in cellular proportions.

**Figure 1.**
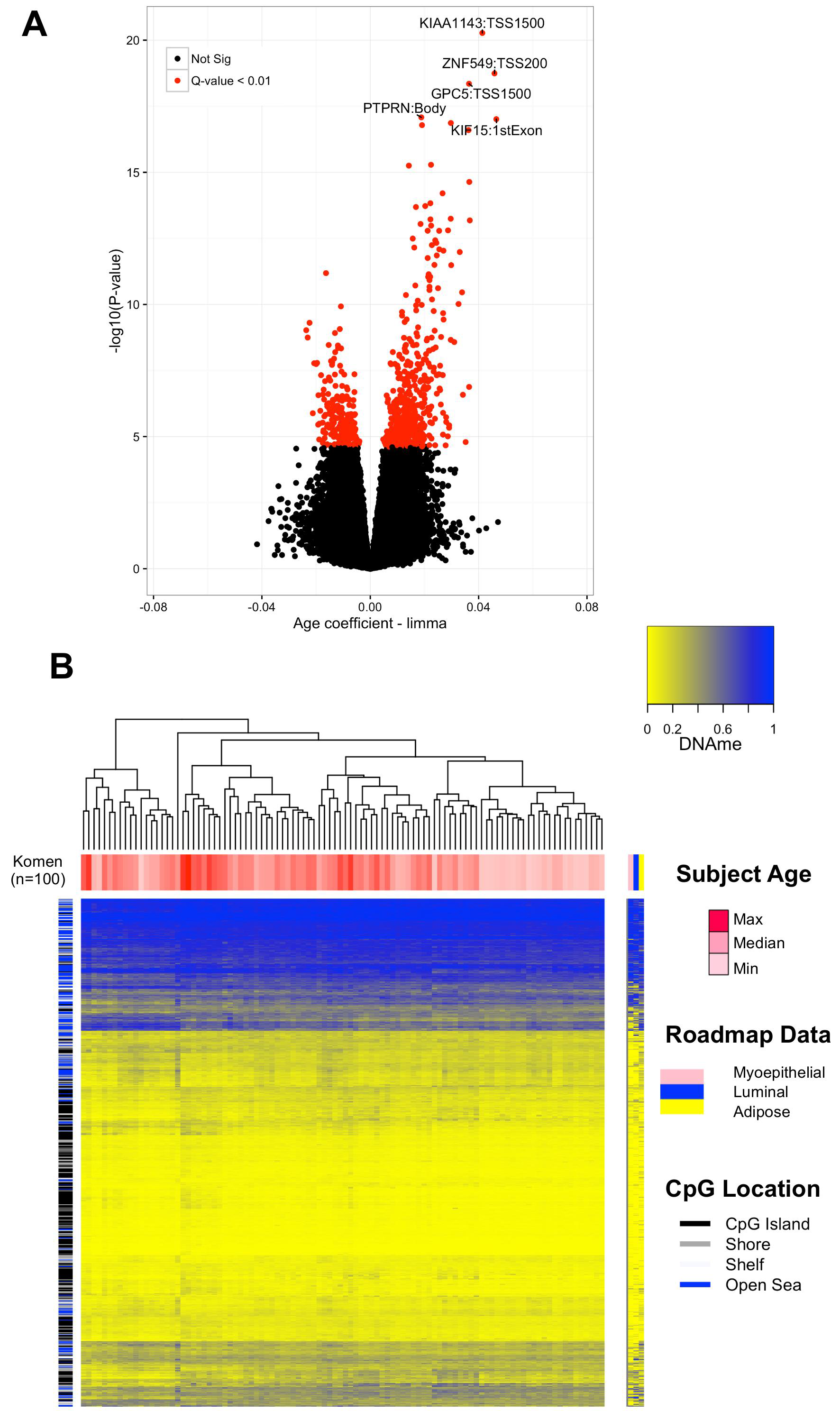
Subject age is strongly associated with DNA methylation in normal breast tissue independent of cell-type. **A.** In the volcano plot, each point represents the associations between DNA methylation and age from cell-type adjusted multivariable limma models at individual CpG sites. Increasing-log10(*P*-value) values on the y-axis show increasing statistical significance and limma effect size on the x-axis positioned away from the zero value reveal the largest DNA methylation changes with age. Significant CpG sites are indicated in red (*Q*-value < 0.01). The gene and gene regions are presented for the 5 CpG sites with the greatest significance. **B.** Unsupervised clustering of DNA methylation values at age-related CpG sites (Komen, n = 100) visualized alongside CpGs measured in specific cell-types form the Roadmap to Epigenomics data set (n = 691 CpG sites). Each column represents a given tissue sample and each CpG in presented in rows.

Visualizations of CpG-specific *P*—values and coefficients from cell-type unadjusted and adjusted models demonstrated that adjustment attenuated both strength and magnitude of CpG-specific associations (**Additional File 4**). Moreover, the number of significant associations (*Q* < 0.01) in the unadjusted limma model for subject age was 4, 099 CpG sites compared with 787 from the adjusted model suggesting that a high number of false-positives are likely to be reported when differences in cell-proportions are not considered (**Additional File 4A-C**). In addition, at the age-related CpG sites (n = 787, *Q* < 0.01) the DNA methylation patterns across purified cell populations of myoepthial cells, luminal cells, and adipocytes are consistent, suggesting that age-related changes may occur largely independent of tissue-type in the normal human breast **Figure 1B**.

There was missing family history data for 10 individuals in the present data set. To explore whether family history was associated with DNA methylation differences we applied the aforementioned limma approach unadjusted and adjusted for cellular proportions (n = 90) and found no significant associations (*Q* > 0.01) between family history and DNA methylation differences after correcting for multiple comparisons (**Additional File 4D**).

### Independent validation of age associated methylation

We next moved to validate our age-related DNA methylation findings in two independent 450K data sets from 97 normal adjacent-to-tumor breast samples (TCGA) and 18 normal breast tissues from disease-free women (NDRI, GSE74214). Subject demographics and characteristics for these two data sets are presented in **Table 2**. In a reference-free cell mixture adjusted limma restricted to the 787 CpG sites identified in the discovery (Komen) population we observed that 548 CpG sites (TCGA, 69.4 %) were differentially methylated in a direction consistent with discovery population at a nominal *P* < 0.05 **Additional File 5A**. Similarly, we observed highly consistent results in the NDRI population (389 out of 787 CpG sites, 49.4%) **Additional File 5B**. Strikingly, there were 345 CpG sites (43.8 %) in the TCGA data set and 109 CpGs (13.9 %) in the smaller NDRI data set that were considered significant at the stringent Bonferroni threshold for multiple comparisons (**Additional File 5A** and **5B**, *P* < 6.4E-05). In both validation cohorts, putative cell mixture proportions were significantly associated subject age (Permutation *P* < 0.05) **Additional File 5C-4D**.

**Table 2.** Independent Population Subject Chacteristics

|  | **Mean (Range)** | **n = 18** |
| --- | --- | --- |
| Age | 49 (13-80) |  |
| BMI | 28.3 (14.59-62.73) |  |

**TCGA**
|  | **Mean (Range)** | **n = 97** |
| --- | --- | --- |
| Age | 57.57 (28-90) |  |
| BMI | Unavailable |  |

While it is appreciated that DNA methylation can modify chromatin structure and distally regulate the transcriptome, its most well defined function is the *cis*-regulation of gene transcription [21]. In the present study, sample-matched RNA-sequencing data was available only for a subset of the subjects from the TCGA dataset (n = 88). Many of the age-related CpG sites that localize to gene regions (n = 630 CpG sites) demonstrated strong associations with gene expression (259 CpG sites at *P* < 0.05, **Additional File 6A**). The direction of the CpG-gene correlations demonstrated a dependency upon genomic context **Additional File 6B**. For example, CpG sites tended to exhibit negative correlations in the promoter region, while there was an even distribution of positive and negative correlations in gene body (that is, intron and exon) regions (**Additional File 6B**).

### Risk factor-associated DNA methylation sites are enriched for regulatory regions

To provide a broader biological interpretation of age-related DNA methylation we next sought to identify enrichment of these genomic locations in gene regulatory regions, such as tissue-specific histone marks and transcription factor binding sites (TFBS). First, we employed the eFORGE tool to identify cell-type specific signals in diverse tissues profiled by the Roadmap to Epigenomics Consortium. We observed robust enrichment of H3K4me1, histone modifications that mark enhancers, in both fetal tissues and mammary epithelial cells (*Q* < 1.9E-37), and modest associations with other histone modifications (that is, H3K4me3, H3K27me3) **Additional File 7**. A Fisher’s exact test confirmed age-related CpGs localize to enhancer elements specifically in mammary myoepithelial cells (H3K4me1, Roadmap) (OR = 2.00 CI (1.73 – 2.33), *P =* 7.1E-20). We next used the genomic coordinates of age-related CpGs as a query set against the background of the 450K array in a Locus Overlap Analysis (LOLA) scanning for enrichments of TFBSs. Since hypermethylation events are likely to be biologically distinct from hypomethylation events at TFBS we stratified our LOLA analysis into a hyper-and a hypomethylation enrichment analysis **Figure 2A-2B**. In the hypermethylation analysis, we observed a striking number of significant enrichments for CpG sites hypermethylated with age (14 TFBS, Q < 0.01) and hypomethylated with age (8 TFBS, Q < 0.01) **Additional File 8A and 8B**. Among the several of the top ranking results presented in **Figure 2A,** MYC and CTCF, which are critical regulators of chromatin architecture were enriched among hypermethylated CpG sites while hypomethylated CpGs localize to binding sites of transcriptional activators c-Fos and Stat-3 [22–25].

**Figure 2.**
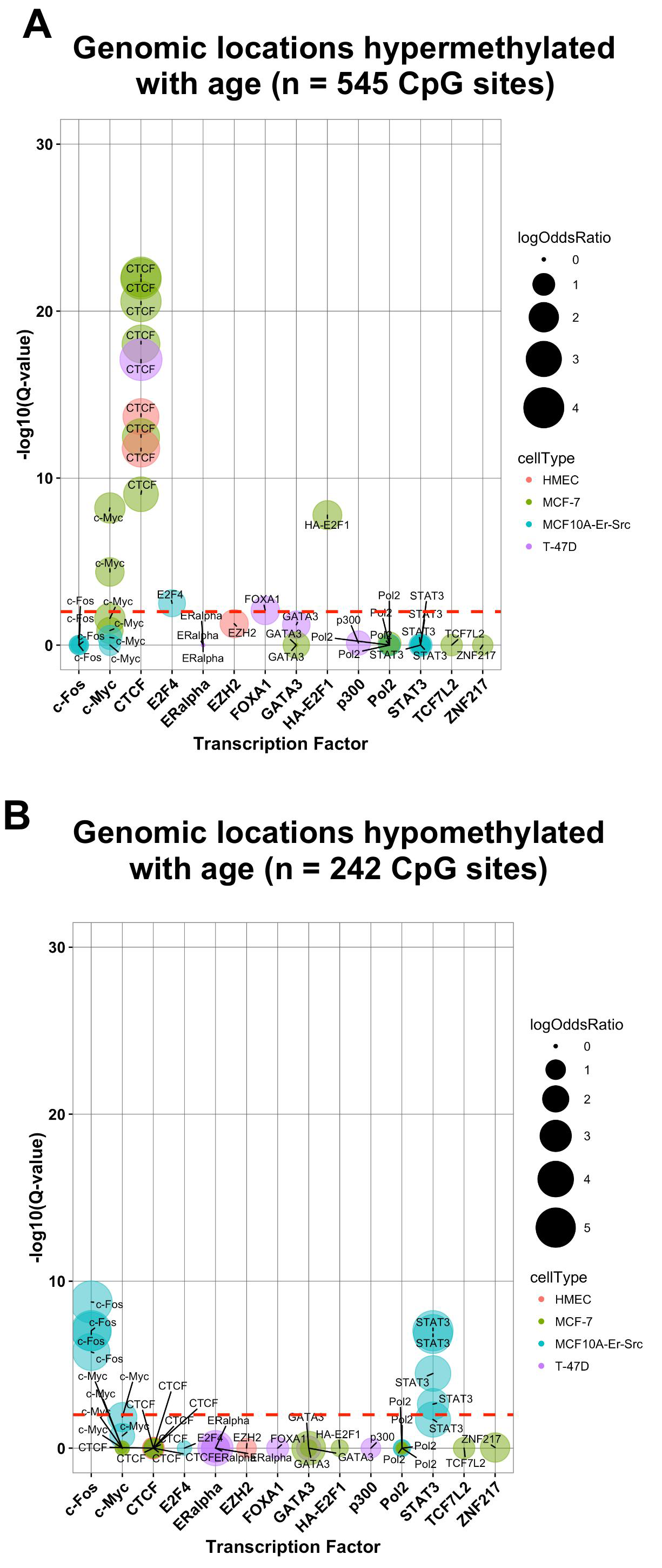
Age-related DNA methylation is enriched for regions of chromatin remodeling and transcriptional control. **A.** Hypermethylated with age CpG sites and **B.** hypomethylated with age CpG sites are highly enriched at the binding sites of transcription factors.

### Accelerated epigenetic aging of human breast tissue

It has been recognized that DNA methylation patterns change in a tissue-specific manner as an individual ages [26]. Previous studies have found that measurements of DNA methylation have the ability to accurately estimate an individual’s age and that observed differences between predicted DNA methylation age (that is, biological age) and chronological age are associated with disease-risk factors [26–29]. Further, it has been observed that DNA methylation age predictions in the human breast demonstrate age acceleration when compared with other tissues suggesting that normal breast tissue tends to age more quickly than other tissues [26].

To examine whether the subject-specific differences between biological and chronological age (that is, age acceleration) are associated with breast cancer risk factors we first calculated DNA methylation age from the 100 Komen normal breast tissues using two distinct epigenetic clocks [26, 29]. Briefly, the “Horvath epigenetic clock” uses elastic net regression to integrate DNA methylation information from 353 CpG sites to generate a multi-tissue age predictor. The second method, “epiTOC”, is an epigenetic clock that incorporates prior biological knowledge into a mathematical model to generate an estimate of mitotic divisions using 385 CpG sites. Notably, there was limited overlap between the 787 age-related CpGs and Horvath (17 CpGs) and EpiTOC (3 CpGs). In analyses with the Horvath clock, we observed a strong positive correlation between chronological age and DNA methylation age of the Komen breast tissues with a Spearman correlation coefficient of 0.95 (*P* = 2.83E-52, **Figure 3A**). In univariate analyses of age acceleration, defined as the residual resulting from regressing DNA methylation age (Horvath clock) on chronological age, and the cancer risk factors listed Table 1 we observed a significant positive association only with race (African-American, *P*= 3.5E-02). Age acceleration was not associated with any other of the evaluated risk factors (*P* > 0.05). In a multivariate model considering all measured cancer risk factors, we found that race was significantly associated with increased epigenetic aging (African-American, *P* = 4.9E-02). In contrast to the Horvath clock, there was no significant correlation between chronological age and epiTOC predicted age (*P* = 7.5E-01, **Figure 3B**). Nonetheless, the epiTOC estimated biological age was also positively associated with race in univariate analyses (African-American *P* = 2.1E-02, Hispanic *P=*2.8E-02) and in multivariate models including all **Table 1** risk factors (African-American, *P* = 2.7E-02 and Hispanic, *P* = 2.7E-02). The remaining breast cancer risk factors were not associated with epiTOC-defined biological aging in either univariate or multivariate models (*P* > 0.05).

**Figure 3.**
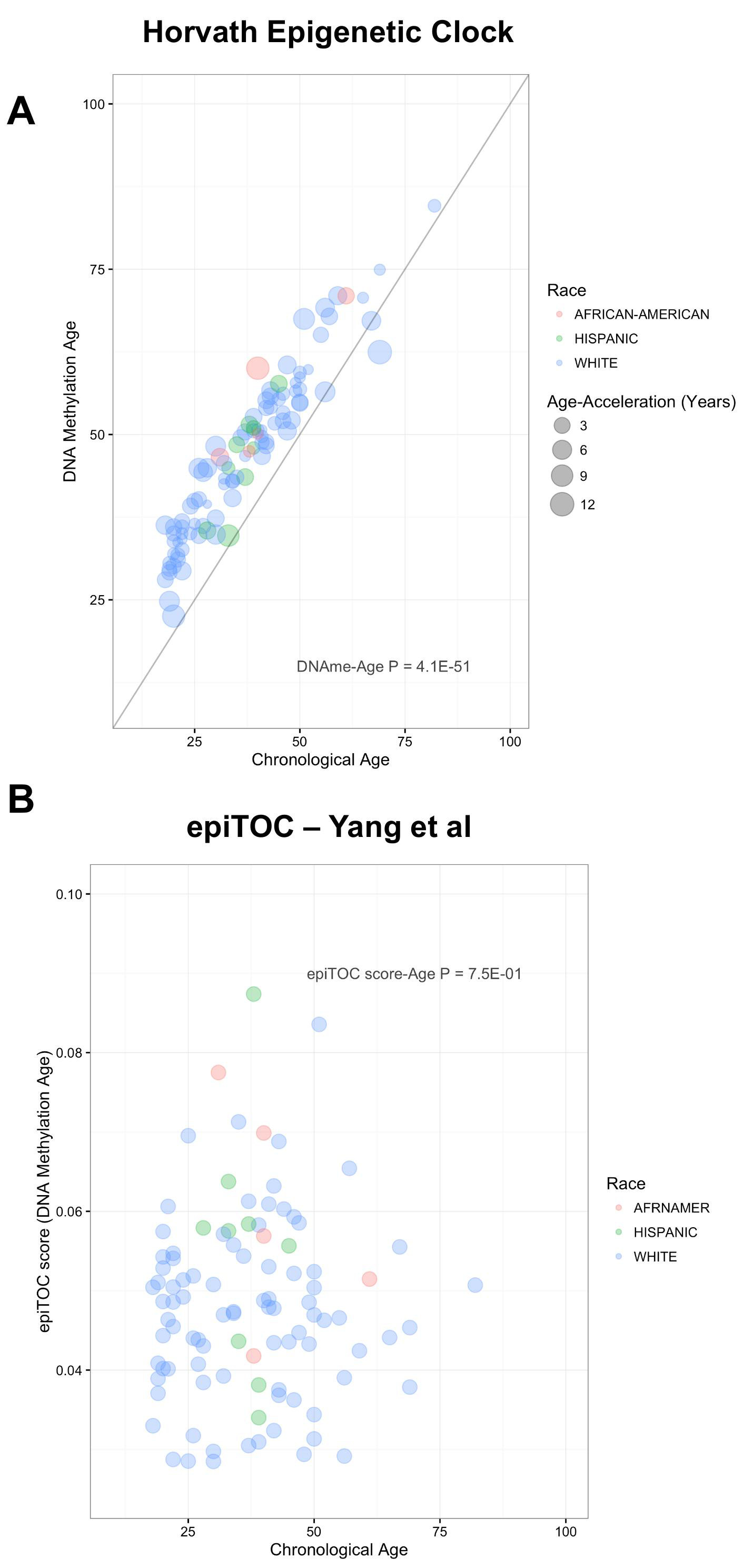
The relation between epigenetic clocks and cancer risk factors. **A.** The Horvath epigenetic clock age in normal breast tissue is highly correlated with subject age (*P* = 2.83E-52). Age acceleration was significantly (*P* < 0.05) larger in African-American women. **B.** DNA methylation age as generated by the epiTOC tool was not correlated with subject age in normal breast tissue (*P* > 0.05). Higher DNA methylation age was associated subject race as breast tissue from African-American and Hispanic women demonstrated increased DNA methylation age (*P* < 0.05).

### Age-related DNA methylation is further deregulated in pre-invasive and invasive breast cancer

To ascertain whether disease risk factor related DNA methylation differences are relevant for the development of cancer we compared DNA methylation in breast tumors with adjacent normal in both pre-invasive and invasive cancer at the 787 age-related CpGs. In pre-invasive lesions (ductal carcinoma in situ, DCIS), there were 268 CpG sites among 775 CpGs available for measure (34.5%) that demonstrated differential methylation between DCIS and normal using limma models adjusted for subject age **Figure 4A** (*P* < 0.05). Importantly, changes at the age-related CpGs were greater (**Additional File 9A and 9B**) and demonstrated stronger associations than a randomly selected set of CpG sites with a similar genomic distribution **Figure 4B** (Kolmogorov-Smirnov test, *P =* 3.0E-03). If the epigenetic defects in age-related DNA methylation are further deregulated in pre-invasive breast cancer it would be expected that progressive changes would occur in invasive breast cancer. To test this end, we assessed differential methylation using limma models adjusted for subject age in the TCGA breast cancer data set. A large proportion of the age-related CpGs exhibited significant differential DNA methylation changes in breast cancer, 642 out of 787 CpGs (81.6%, *P <*0.05) **Figure 4C**. Again, we found that the age-related changes demonstrated greater DNA methylation differences (**Additional File 9C and 9D**) and stronger associations than a randomly selected set of CpGs (Kolmogorov-Smirnov test, *P =* 1.1E-13) **Figure 4D.**

**Figure 4.**
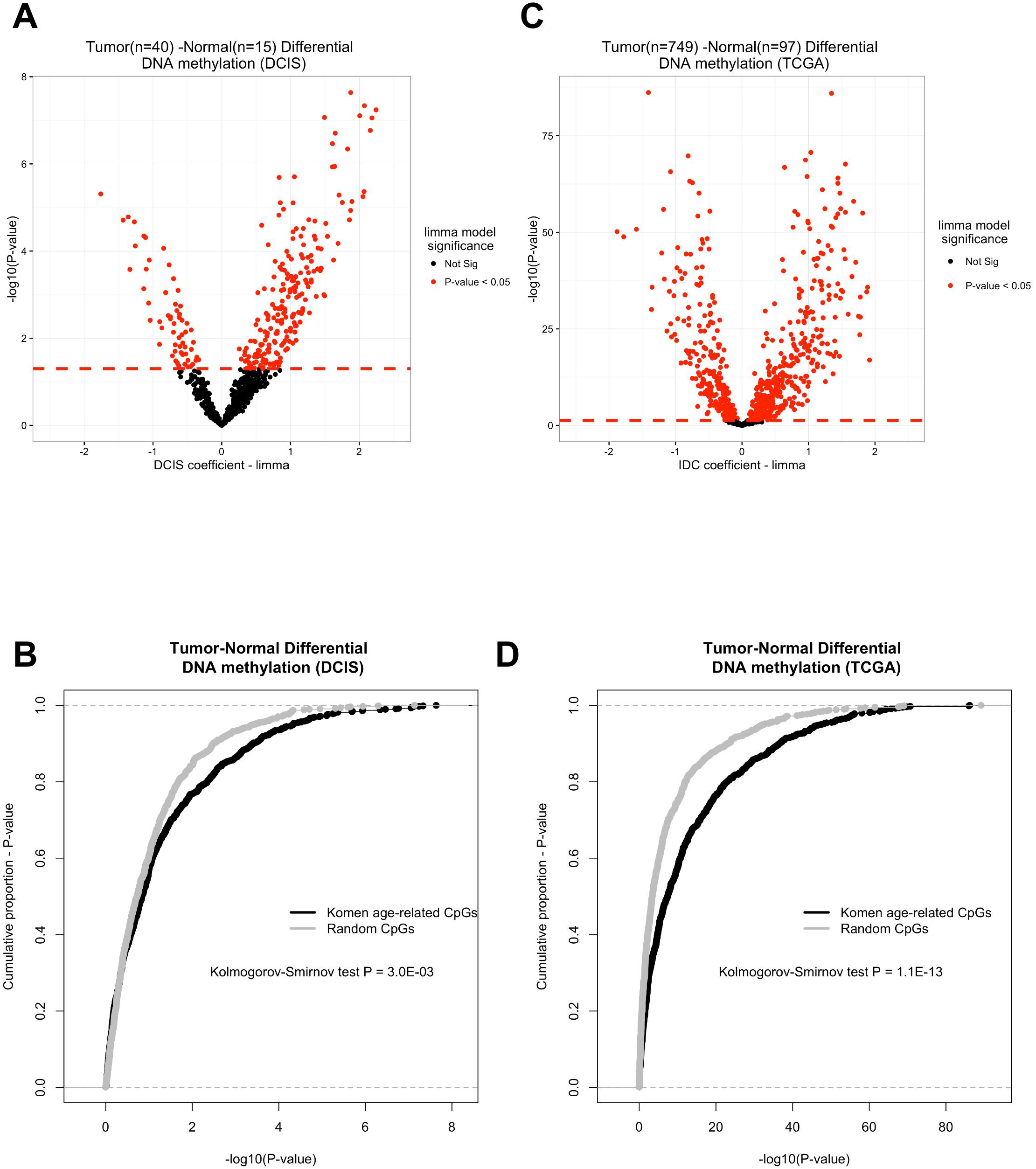
DNA methylation differences between tumor and normal breast tissue at age-related CpG sites in both **A-B.** ductal carcinoma *in situ* (DCIS) and **C-D.** invasive breast cancer.

## Discussion

In this study, we identified perturbations in the normal breast epigenome that may contribute to age-related increases in breast cancer risk. Age is the strongest demographic risk factor for breast cancer and is robustly associated with DNA methylation changes. Emerging literature has demonstrated that aging exerts it profound effects on the epigenome through a lifetime accumulation of environmental exposures that interfere with the placement or removal of methyl groups [12, 13, 30, 31]. Here, we have described that the consistent changes in breast DNA methylation are not randomly distributed throughout the genome. Instead, age-related DNA hypermethylation events are enriched for breast epithelial-specific enhancer regions and the binding sites of chromatin remodelers while hypomethylation was noted at transcriptional activators. The enrichment of modifications at critical regulators of cellular phenotype provide novel insights into how cell type-specific epigenetic states change over time and may predispose cells to neoplastic transformation. Our analysis revealed that further DNA methylation alterations to these genomic regions in pre-invasive and invasive disease may contribute to the restriction of cellular differentiation and disruption of transcriptional control observed in cancerous lesions.

The ability to produce reliable biological age predictions for an individual as well as specific tissues holds promise for monitoring health, predicting disease risk, and providing insights about modifiable lifestyle factors that promote healthy aging. Indeed, discrepancies found between chronological and biological age may suggest deregulation in DNA methylation marks and indicate increased disease risk. Horvath *et al.* demonstrated this phenomenon of age-acceleration in a recent publication where researchers found that the epigenetic age of liver was increased by 2.7 years for every 10 units of body mass index [27]. Using 450K methylation arrays we have applied the Horvath epigenetic clock algorithm and epiTOC tool to 100 normal tissue samples to determine the DNA methylation age of each of these tissues. The association between age acceleration and race produce novel hypotheses for risk given the observation that African-American women tend to be diagnosed with breast cancer at an earlier age [32]. In future studies, the ability to accurately assess biological age in breast tissue samples from larger longitudinal studies with a greater number of African-American and Hispanic women may aid researchers in the determination of factors that aim to assess and prevent disease.

While our findings provide strong evidence for a link between epigenetic deregulation and the two processes of aging and cancer our study holds a few limitations. For example, although the RefFreeEWAS method effectively accounts for the largest sources of variation in the DNA methylation data set the method is unable to discern in which particular cell-types the epigenetic changes occur. That said, the robustness of cell-type independent observation across multiple populations and progressive alterations in cancer gives us confidence that a subset of the epigenetic defects may be important in carcinogenesis. To this end, future prospective studies are needed to investigate the relation of DNA methylation in normal tissue with risk of developing breast cancer. Research aimed at early detection and disease prevention would serve to relieve patient morbidity and reduce extra cost to the healthcare system. In summary, we have shown that epigenetic differences are strongly associated with aging and these differences may reflect epigenetic defects that predispose women at an older age to an increased risk of breast cancer.

## Conclusions

Epidemiological studies have firmly established factors of personal choice as well as factors beyond personal choice that alter risk of breast cancer. Established risk factors for breast cancer include age, reproductive and family history, as well as body mass index (BMI) [5, 33]. Indeed, modeled breast cancer risk factors have been shown to account for approximately half of breast cancer cases [34, 35]. However, the biological mechanisms by which specific risk factors impact disease risk are not well understood. In this study population, we did not observe significant associations between BMI or parity and genome-wide DNA methylation. However, we observed consistent cell-type independent age-related DNA methylation in multiple populations of normal breast tissue. The genomic locations of age-related DNA methylation were more likely to be found in gene regulatory elements of breast epithelial cells suggesting a loss of cellular state control as an individual ages. Further, we demonstrate additional support for a link between age-related DNA methylation and cancer, as age-related CpG sites were more likely to exhibit greater alterations in both pre-invasive and invasive breast cancer. Together, our research suggests that DNA methylation changes in aging shifts the epigenetic state toward a compromised molecular phenotype creating a novel link between risk factors and potential disease origins in breast cancer.

## Methods

### Study population

The discovery population consisted of 100 cancer-free women who donated breast tissue biopsy specimens to the Susan G. Komen Tissue Bank after providing written informed consent. We selected biospecimens from women with a biopsy that scored for a high proportion of epithelial cells as determined by the Susan G. Komen Tissue Bank study pathologist (n=100) [36]. The sample population was selected for an approximately equal distribution of parous and nulliparous women, and to include a wide age range of subjects. Subject demographic and breast cancer risk factors were collected from tissue donors using a questionnaire administered by the Susan G. Komen Tissue Bank. Family history of cancer was defined by whether or not the donor had at least one first-degree blood relative (i.e., mother or sister) diagnosed with breast cancer.

### DNA methylation quantification and normalization

Fresh frozen tissue samples were manually dissected and DNA was extracted using Qiagen DNeasy Blood and Tissue Kit according to manufacturer’s protocol (Qiagen, Valencia, CA, USA). DNA was quantified using a Qubit fluorometer and 1ug of DNA was then bisulfite modified using the EZ DNA methylation kit (Zymo research, Orange, CA, USA) according to manufacturer’s recommended protocol. The resulting material was used as input for the hybridization on the Infinium HumanMethylation450 BeadChip (Illumina, San Diego, CA, USA). Samples were randomized to plates and subjected to epigenome-wide DNA methylation assessment. The methylation status for each CpG locus was calculated as the ratio of fluorescent signals (β = Max (M, 0) / [Max(M, 0) +Max(U, 0)+100]), ranging from 0 (non-methylated) to 1 (completely methylated), using average probe intensity for the methylated (M) and unmethylated (U) alleles. Normalization and background correction of raw signals was performed using the *FunNorm* procedure available in the R/Bioconductor package *minfi*(version 1.10.2) [12]. Illumina probe-type normalization was carried out with beta-mixture quantile normalization (BMIQ) [37]. Prior to analysis we removed CpG sites on sex chromosomes as well as those corresponding to probes previously identified as cross-reactive or containing SNPs, resulting in 390,292 CpGs remaining for analysis [38].

### Validation in Independent Populations and The Cancer Genome Atlas

Independent breast tissue samples were available from the National Disease Research Interchange (NDRI, GSE74214, n = 18) and The Cancer Genome Atlas Database (TCGA, n = 97) [39]. Raw intensity data (IDAT) files were available for both studies and DNA methylation data was processed and normalized using the same methods described above. Likewise, raw DNA methylation IDAT files were accessed and processed using the same methods outlined above for both ductal carcinoma *in situ* (n=55, GSE66313) and invasive ductal carcinoma (n=749, TCGA) to compare DNA methylation differences between normal-adjacent tissue and pre-invasive or invasive lesions [9].

### Statistical Analysis

All data analysis was conducted in R version 3.3.1.

*Cell-Mixture Deconvolution*. Adjustment for variation in cellular proportions can be achieved in the absence of referent DNA methylation dataset from characterized cell types [15, 20]. To perform a reference-free epigenome-wide association study (EWAS) we used the R package RefFreeEWAS to deconvolute the cellular populations present in the tissue biopsy samples using DNA methylation data as detailed previously in Houseman *et al*. [20]. Briefly, this method seeks to represent the largest axes of variation in the DNA methylation data set and decomposes the DNA methylation data for a sample of heterogeneous cell populations into its constituent methylomes. As a convex variant of non-negative matrix factorization, the RefFreeEWAS method is similar to approaches used to deconvolute gene expression levels in heterogeneous tumor tissues [40, 41]. In the present study, we selected the 10,000 most variable CpGs in each data set and used a bootstrap technique (specifically sampled the specimens with replacement 1,000 times) to estimate the optimal number of putative cell types (K). The optimal number of cell-types defined in each data set were: K = 6 (Komen), K = 10 (TCGA adjacent normal), and K = 2 (NDRI normal breast). The discrepancy in estimated cell-types for each population can be explained in part by sample size (i.e., small for NDRI population) and potential epigenomic field defects in normal-adjacent to tumor tissue (i.e., TCGA).

*Analysis of CpG-Specific Associations*. We used a multivariable limma procedure as described in the R/bioconductor library *limma* [42] to model CpG-specific associations between logit-transformed beta values (i.e., *M*-values) and breast cancer risk factors (e.g., age, body mass index, parity). Genome-wide significance was determined by taking into account the false discovery rate with a threshold of statistical significance set at *Q* = 0.01. We ran separate multivariate limma models both unadjusted and adjusted for putative cell proportions to assess the impact of cell proportion differences on significant associations and effect-size estimates. To identify loci that may be most confounded by differences in cell-type we calculated the difference in the effect size estimates (i.e., delta coefficient value) between the cell-type unadjusted and adjusted models.

*Associations with Metadata*. To test the associations between putative cellular proportions and subject metadata (e.g., age) we applied the methods described in Houseman *et al*. to fit a quasi-binomial model for each putative cell-type across the data set [20]. More specifically, for each estimated value of K (that is, total number of cell-types), we generated a model for each cell-type (1 to K) and used the minimum *P*-value. We then computed the permutation distribution of these minimum *P*-values (spanning all potential values of K).

*Genomic Region Enrichment*. To assess the enrichment of risk factor-related CpG sites at cell type-specific histone modifications we used the eFORGEv1.2 tool with the selected option of all H3 marks measured for the consolidated Roadmap to Epigenomics data set [43]. To examine whether risk factor-related CpGs were associated with transcription factor binding sites in ENCODE data we used the Locus Overlap Analysis (LOLA) software [44]. In this analysis, our query input set of genomic regions to be tested for enrichment were the genomic locations of the risk factor-related CpG sites (*Q* < 0.01) and the background set was the genomic locations of the 390,262 CpGs used in the entire analysis. For the LOLA analysis, the ENCODE transcription factor binding sites included 42 different ChIP-seq experiments.

*Epigenetic Clock Analysis*. DNA methylation age (biological age) for the Komen breast tissues was calculated using the Horvath and EpiTOC methods [26, 29].

## Availability of data and materials

The DNA methylation microarray data from healthy Komen Tissue Bank donors that support the findings of this paper have been deposited in the Gene Expression Omnibus with the accession codes GSE88883 (http://www.ncbi.nlm.nih.gov/geo/). The DNA methylation microarray data from an independent population of non-disease breast tissue (NDRI) have been deposited in the Gene Expression Omnibus with the accession codes (GSE74214). Level 1 IDAT and Level 3 normalized RNASeqV2 rsem.genes.normalized_results were downloaded from The Cancer Genome Atlas breast cancer project (TCGA, http://cancergenome.nih.gov). R code used for analyses presented in this manuscript has been deposited in the “Normal-Breast-Methylation” repository on github (https://github.com/Christensen-Lab-Dartmouth).

## List of abbreviations

5mC: 5-methylcytosine
CpG: Cytosine-guanine dinucleotide
TCGA: The Cancer Genome Atlas
NDRI: National Disease Research Interchange
TFBS: transcription factor binding site
PCGT: Polycomb group protein target
TSS: transcriptional start site
KEGG, GEO: Gene Expression Omnibus
epiTOC: Epigenetic Timer of Cancer
IDAT: intensity data file
limma: linear models for microarray data
ENCODE: Encyclopedia of DNA Elements
LOLA: locus overlap analysis
BMI: body mass index
EWAS: Epigenome-wide association study
RefFreeEWAS: Reference-Free DNA Methylation Mixture Deconvolution Epigenome-wide association study

## Competing interests

The authors declare that they have no competing interests.

## Funding

The research reported in this publication was supported the Center for Molecular Epidemiology COBRE program with grant funds from the National Institute of General Medical Sciences (NIGMS) of the National Institutes of Health under award number P20 GM104416 (PI: Margaret R. Karagas). This work was supported by the National Institutes of Health grant numbers R01DE022772 to BCC, R01MH094609 to EAH.

## Author’s contributions

KCJ conceived and designed the approach, carried out laboratory experiments, performed statistical analyses, interpreted the results, wrote and revised the manuscript. EAH interpreted the results, generated statistical framework, and revised the manuscript.

JEK carried out laboratory experiments and revised the manuscript.

BCC conceived and designed the approach, oversaw project development, interpreted the results, and revised the manuscript. All authors have read and approved the final manuscript.

## Acknowledgements

Samples from the Susan G. Komen Tissue Bank at the Indiana University Simon Cancer Center were used in this study. We thank contributors, including Indiana University who collected samples used in this study, as well as donors and their families, whose help and participation made this work possible.

## Additional Files

**Additional File 1:** Analytic framework for reference-free epigenome-wide association study between DNA methylation and breast cancer risk factors.

**Additional File 2:** Estimation of cellular proportions and its association with subject covariates. **A.** Hierarchal clustering and heatmap representation of cellular proportions of putative cell-types (K = 6) in Komen normal breast tissue (n = 100). **B.** Metadata associations with cellular proportions when K is estimated over a range of cell types. Permutation *P*-values presented adjacent to the colored line representing each covariate (e.g., red for age, permutation *P* = 2.0E-03).

**Additional File 3:** Age-related CpGs and genomic annotation

**Additional File 4:** Volcano plots representing both cell-type proportions adjusted and unadjusted limma models for each covariate evaluated in the present study. In each cell-type adjusted volcano plot (right panels) the intensity of blue and red colored points indicate shift in the effect size of the limma coefficient estimate between adjusted and unadjusted models. That is, gray points in the right panels indicate CpG sites that are not impacted by differences in cellular proportions across **A.** subject age (n =100) **B.** subject BMI (n = 100) **C.** parity status (n =100) **D.** and family history of disease (n = 90).

**Additional File 5:** Age-related DNA methylation in the human normal breast validates in **A.** adjacent-to-tumor normal breast from The Cancer Genome Atlas (TCGA, n = 97) population and **B.** the National Disease Research Interchange (NDRI, n = 18) normal breast tissue population. Volcano plots indicate CpG-specific associations between DNA methylation and subject age. Permutation testing of subject covariate data across estimated cell-types (K) in **C.** TCGA population and **D.** NDRI population.

**Additional File 6:** Age-related CpG sites are associated with gene transcription. **A.** Distribution of *P*-values for CpG-gene expression correlations **B.** Genomic-context dependency between DNA methylation and gene expression. Gene names for the 20 CpG-gene regions with the strongest associations are presented alongside its respective coefficient-*P*-value bubble.

**Additional File 7:** Complete results from eFORGE analysis of age-related CpGs (n=787).

**Additional File 8:** Complete results from LOLA analysis of age-related CpGs (n=787).

**Additional File 9: A-B.** DNA methylation differences between DCIS and normal adjacent in limma coefficients (that is, effect size) for age-related (n =787) and randomly selected loci (n = 787) **C-D**. DNA methylation differences between invasive breast cancer and normal adjacent in limma coefficients (that is, effect size) for age-related (n = 787) and randomly selected loci (n = 787).

